# Heart development in the lizards (Varanidae) with the greatest extent of ventricular septation

**DOI:** 10.1101/563767

**Authors:** Jermo Hanemaaijer, Martina Gregorovicova, Jan M. Nielsen, Antoon FM Moorman, Tobias Wang, R. Nils Planken, Vincent M Christoffels, David Sedmera, Bjarke Jensen

**Author notes:** These authors contributed equally (Jermo Hanemaaijer, Martina Gregorovicova). corresponding author: Bjarke Jensen, Ph.D., Amsterdam UMC, Department of Medical Biology, Room L2-106, Meibergdreef 15, 1105AZ Amsterdam, The Netherlands, Fax: not available.

## Abstract

Among lizards, only monitor lizards (Varanidae) have a functionally divided cardiac ventricle. This enables them to sustain higher systemic blood pressures and higher metabolic rates than other reptiles of similar size. The division results from the concerted action of three partial septa, which may have homology to the full ventricular septum of mammals and archosaurs. Homology, however has only been inferred from anatomical comparisons of hearts of adult monitors whereas gene expression during heart development has not been studied. We show in developing monitors that the partial septa that separate the left and right ventricle, the ‘muscular ridge’ and ‘bulbuslamelle’, express the evolutionary conserved transcription factors *Tbx5, Irx1* and *Irx2*, orthologues of which mark the full ventricular septum. Compaction of embryonic trabeculae contributes to the formation of these septa. The septa are positioned, however, to the right of the atrioventricular junction and they do not partake in the separation of incoming atrial blood streams. Instead, the ‘vertical septum’ within the left ventricle separates the atrial blood streams. It expresses *Tbx3* and *Tbx5*, which orchestrate the formation of the electrical conduction axis of the full ventricular septum. These patterns of expression are more pronounced in monitors than in other lizards, and are associated with a deep electrical activation near the vertical septum, contrasting the primitive base-to-apex activation of other lizards. We conclude that current concepts of ventricular septum formation apply well to the monitor septa and that there is evolutionary conservation of ventricular septum formation among amniote vertebrates.

## Introduction

The evolution of land-living vertebrates was facilitated by breathing of air with lungs (Perry & Sander, 2004). A dedicated vascular bed, the pulmonary circulation, enables the perfusion of the lungs by blood and the oxygenated blood of this circulation returns to the left side of the heart (Burggren & Johansen, 1986). The oxygen-rich blood from the lungs, however, may mix with oxygen-poor blood returning from the systemic circulation if septal structures are absent in the ventricle of the heart. In amphibians, which include the earliest land-living vertebrates, the ventricle is without a septum whereas in mammals and in archosaurs (crocodylians and birds) the ventricle is divided into the left and right ventricle by a full ventricular septum (R. E. Poelmann et al., 2014). Mammals evolved from reptile-like ancestors and archosaurs evolved within reptiles (Laurin & Reisz, 1995), and among the extant non-archosaur reptiles there is substantial variation in the degree of ventricular septation (B. Jensen, Moorman, & Wang, 2014).

The single cardiac ventricle of extant non-archosaur reptiles is partially divided by three septa named the “vertical septum”, the “muscular ridge” and the “bulbuslamelle” The ‘vertical septum’ is a prominent sheet of trabecular myocardium positioned beneath the atrioventricular valve (Fig. 1). It separates the oxygen-rich blood from oxygen-poor blood coming from the right atrium and its function therefore resembles that of the ventricular septum of mammals and archosaurs (B. Jensen et al., 2014). The muscular ridge is the most prominent septal structure, but is not involved in the separation of the atrial inflows because it is positioned to the right of the atrioventricular canal (R. E. Poelmann et al.). The ventricular septum of mammals and archosaurs, in contrast, is positioned immediately below the atrioventricular canal (Cook et al.; Greil, 1903; Webb, Heatwole, & Bavay) (Fig. 1). During ejection, the muscular ridge, however, forms an important boundary between the pulmonary compartment, the cavum pulmonale, and the cavum venosum to separate blood flows into the aortae and the pulmonary artery (Fig. 1). The third septum, the ‘bulbuslamelle’, is positioned opposite to the muscular ridge and presses against the free edge of the muscular ridge during contraction. It has been shown experimentally that this coming together of the two septa prevents mixing of oxygen-rich blood and oxygen-poor blood during cardiac contraction (White, 1959). The full ventricular septum of mammals, birds and crocodylians has a pronounced left-right gradient of expression of the transcription factor *Tbx5* (B. Jensen et al.; Koshiba-Takeuchi et al., 2009; R.E. Poelmann et al., 2014). Analyses of *Tbx5* gradients suggested that the vertical septum of a turtle (*Trachemys*), but not the vertical septum of a lizard (*Norops*), shows homology to the full ventricular septum (Koshiba-Takeuchi et al.). Studies on other species of turtles and squamate reptiles, however, show that both the muscular ridge and the vertical septum display a gradient of *Tbx5*expression (R.E. Poelmann et al., 2014). The seemingly conflicting results in non-crocodylian reptiles could be resolved if the expression of *Tbx5* identifies more than one septal structure in animals without a full ventricular septum.

**Fig. 1.**
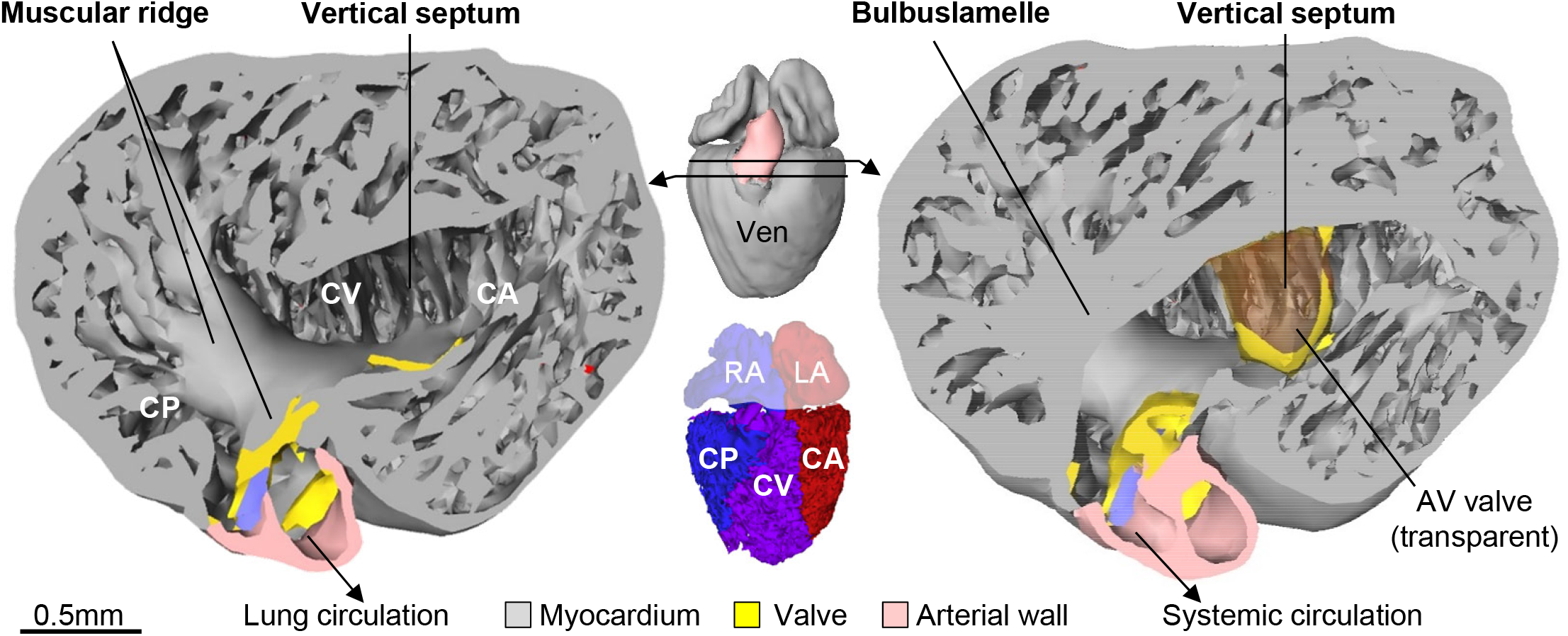
Septa and sub-compartments in a typical lizard ventricle. The luminal side of the lizard ventricle, looking towards the ventricular apex, shown from two transverse cuts made in the ventricular base. Images from 3D model of the heart of an adult anole lizard, from (B. Jensen et al., 2014). There are three septa (muscular ridge, bulbuslamelle, vertical septum), none of which fully separates the ventricular cavity. Accordingly, the division into three sub-compartments is not based on strict boundaries, but inferred from anatomy and blood flow (B. Jensen et al., 2014): cavum arteriosum (CA, red in insert), cavum venosum (CV, purple in insert), and cavum pulmonale (CP, blue in insert). Notice the vertical septum is beneath the atrioventricular valve (AV valve, made transparent). LA, left atrium; RA, right atrium; Ven, ventricle.

Because the cardiac anatomy of the last common ancestor is likely to remain unknown, heart fossilization is extremely rare (Maldanis et al., 2016), we can only provide insight into the evolution of the ventricular septum through studies of the cardiac anatomy in extant reptiles. Among these, the monitor lizards (Varanidae) have ventricular septa that are more developed than in any other lizards (B. Jensen et al., 2014; Webb et al., 1971). Further, monitors are exceptional among lizards because their cardiac ventricle is functionally divided, such that the left side of the ventricle (cavum arteriosum and venosum) generates high systemic blood pressure whilst the cavum pulmonale supplies the lungs at much lower pressure (Burggren & Johansen, 1982). These mammal-like blood pressures support the metabolic rates that are higher than in other reptiles (Thompson & Withers, 1997) and the sequencing of the Komodo dragon genome suggests that many mitochondrial genes of monitors have undergone positive selection (Lind et al., 2019). The greater state of development of the septa could associate with more pronounced patterns of gene expression and the monitor ventricle may then be the most suitable model to test the evolutionary origin of the full ventricular septum.

To investigate whether the ventricle of monitors is divided by structures that are homologous to the full ventricular septum of mammals and archosaurs, we studied heart development and function in monitors (*Varanus acanthurus, V. exanthematicus, V. indicus*, and *V. salvator*) and compared them to phylogenetically distant lizards with typical lizard hearts, the Brown Anole (*Norops sagrei*) and the Leopard Gecko (*Eublepharis macularius*). To assess homology between amniote cardiac structures, we studied the expression of anole orthologues of genes associated with the formation of the ventricular septum of mammals and birds, specifically *Irx1, Irx2, Tbx3, Tbx5* and *Myh6* (Aanhaanen et al., 2010; Boukens & Christoffels, 2012; Bruneau et al., 1999; Christoffels, Keijser, Houweling, Clout, & Moorman, 2000; de Groot et al., 1987; Hoogaars et al., 2004; A. F. M. Moorman, de Jong, Denyn, & Lamers, 1998; Yamada, Revelli, Eichele, Barron, & Schwartz, 2000). We then related the degree of septation to electrical function. In mammals, the activating current spreads from the atrioventricular node to the atrioventricular bundle that resides in the crest of the ventricular septum (Durrer et al., 1970; van Rijen et al., 2001). This mode of activation reveals itself on the epicardial surface as an early activation near the ventricular apex (Chuck, Meyers, France, Creazzo, & Morley, 2004; Reckova et al., 2003; Rentschler et al., 2001; Sankova et al., 2012). Electrical activation is much the same in crocodylians, which have a full ventricular septum, whereas activation of the undivided ventricle in lizards and snakes proceeds in a primitive pattern from base to apex (Christian & Grigg, 1999; Gregorovicova, Sedmera, & Jensen, 2018; B. Jensen et al., 2018; B. Jensen et al., 2012). Possibly, the advanced state of ventricular septation in monitor lizards could impact on electrical propagation.

We show in monitors that the ventricular septa express orthologues of genes associated with the formation of the ventricular septum of mammals and birds and that the pattern of electrical activation is specialized compared to other squamate reptiles. The ventricle of monitors is seemingly an extreme variation of the common design to the non-crocodylian reptilian ventricle.

## Materials and Methods

### Animals

An adult Savannah Monitor (*Varanus exanthematicus*, Bosc 1792) was used for echocardiography (ca. 2kg). The heart of one 11-year old Water Monitor (*V. salvator*, 10kg, used for MRI) was donated to us (BJ) after the animal died from senescence. From six fertilized eggs of Ridge-tailed Monitor (*Varanus acanthurus*, Boulenger 1885) that were incubated at 30°C and 80-90% humidity, we investigated hearts isolated on the following days post oviposition (dpo): 15, 21, 25, 32, 39, and 46. From 11 fertilized eggs of Leopard Gecko (*Eublepharis macularius*, Blyth 1854) that were incubated similarly, hearts were isolated on the following days: 7, 11, 15 (N=2), 18, 26 (N=2), 32, 40, 47 and 53. We further made use of a previously published developmental series of the Mangrove Monitor (*Varanus indicus*, Daudin 1802) (Gregorovicova, Zahradnicek, Tucker, Velensky, & Horacek, 2012). The hearts of this series were stained with immunohistochemistry for heart muscle sarcomeric proteins only, as initial tests showed that the tissue preservation was not good enough for the detection of most antigens and gene transcripts by *in situ* hybridization. Of the Brown Anole (*Norops sagrei*, Duméril and Bibron 1837), we investigated embryos of Sanger stages (Sanger, Losos, & Gibson-Brown, 2008) 7 (N=1), 9 (N=1), 11 (N=2), 16 (N=1), and one adult, approximately 1 year old (the handling of the adult Brown Anole complied with Dutch national and institutional guidelines (Amsterdam UMC, the Netherlands) and with the Institutional Animal Care and Use Committee of the University ratified approval registered as “DAE101617”. All data was generated within the time limits of “DAE101617”). The heart of *Chelodina mccordi, Chelydra serpentine* and *Cyclodomorphus gerrardii* came from animals that were bought commercially, and housed and euthanized at Aarhus University in accordance with national and institutional guidelines.

### Echocardiography

One adult Savannah Monitor (*V. exanthematicus*) was anaesthetized with isoflurane and intubated for manual ventilation with room air containing 1.5 % isoflurane. Images were obtained at room temperature while the lizard was placed on its back on a heating pad set at 25° C. We used a Vivid 7 echocardiographic system (GE Healthcare, USA) with an 11 MHz phased array pediatric transducer operating at a frame rate of 60 Hz. Two-dimensional images were recorded in the three principal planes of the body (sagittal, transverse and frontal) and stored for off-line evaluation on a workstation (EchoPac Dimension 06, GE Healthcare, USA).

### Optical mapping

Optical mapping was performed on *V. acanthurus* and *Eublepharis macularius*. Shell and membranes were removed and the embryos were placed in dish containing ice-cold reptilian Ringer solution (specific Ringer solutions for *Norops* (adopted from (B. Jensen et al., 2012)); in mmol/l: NaCl 95, Tris 5, NaH_2_PO_4_ 1, KCl 2.5, MgSO_4_ 1, CaCl_2_ 1.5, Glucose 5, pH adjusted to 7.5 with HCl). The heart and adjacent posterior body wall structures were isolated and stained in 2.5 mmol/l di-4-ANEPPS (Invitrogen) for 10 min. Contractions were inhibited with cytochalasin D (0.2 μM, Sigma) and the torso was pinned to the bottom of the dish containing oxygenated Ringer at 28 °C. Imaging was performed from both the ventral and dorsal surfaces at 0.25-1 kHz. Ventricular pacing was performed in the three oldest specimens of both species using a platinum electrode at 125% of the intrinsic rate and twice the diastolic threshold (Sedmera et al., 2003). Data acquisition and analysis were performed using the Ultima L high-speed camera and bundled software (Sankova et al., 2012). Epicardial activation maps were then constructed separately for the ventral and dorsal (as well as lateral, when available) ventricular surfaces in sinus and stimulated rhythm.

### Fixation, *in situ* hybridization, and immunohistochemistry

For *V. acanthurus*, the hearts, and surrounding tissues, were fixed for one day in freshly made 4% paraformaldehyde in PBS (0.9% NaCl) and then washed twice in PBS before finally stored in 70% ethanol. For *in-situ* hybridization we used previously used probes for *Norops* mRNA (*Tbx3, Tbx5, tnnt2, Myh6;(B*. Jensen et al.; B. Jensen et al., 2012)) and new probes based on the following coordinates using UCSC Genome Browser on Lizard May 2010 (Broad AnoCar2.0/anoCar2) Assembly; *Irx1* (chrUn_GL343292:777,311-778,155), *Irx2* (chrUn_GL343292:1,436,684-1,440,677). Tests with *Norops Bmp2* and *Gja5* (Cx40) (also from (B. Jensen et al., 2012)) did not show specific staining in *Varanus. In-situ* hybridizations were performed as described previously (A. F. Moorman, Houweling, de Boer, & Christoffels, 2001) and done on series of sections of 8, 10, or 12 μm thickness that contained the entire heart of specimens of all *V. acanthurus*. Sections of mouse and chicken hearts with detection of *Irx1* and *Irx2* were adapted from (Christoffels, Habets, et al., 2000). For immunohistochemistry, sections of the embryonic hearts of *V. indicus* were cooked under pressure for 5 min in antigen retrieval solution (H-3300, Vector). Myocardium was labelled with a mouse polyclonal antibody to cardiac troponin I (Millipore, dilution 1:250, RRID:AB_2256304). The antibody was detected with a fluorescently labeled secondary donkey-anti-mouse antibody (Invitrogen, dilution 1:200) with Alexa Fluor 488. Subsequently, sections were mounted using glycerol-PBS (1:1) containing SYTOX^®^ Blue (Molecular Probes, dilution 1:40.000) for nuclear staining.

### Magnetic Resonance Imaging (MRI)

For the MRI acquisition of the heart of the *V. salvator*, a 3.0T MR scanner (Ingenia; Philips Medical Systems, Best, The Netherlands) was used. For signal reception, a 32-channel head coil was used. For the 3D MRI acquisition, a T1 weighted turbo-field echo (TFE) pulse sequence was used with the following imaging parameters: TR / TE = 6.7 / 2.9 ms, FA = 20 deg, acquired voxel size = 0.5×0.5×0.5 mm, NSA = 2. Total scan duration was 7 min 23 s. The presented image is a thin MinIP (minimal intensity projection) MR image reconstruction (5 mm slice thickness).

### 3D reconstructions

Reconstructions were made in Amira software (FEI) version 5.5 as described previously (Soufan et al., 2003). In the few cases where a section was damaged (less than 10% of all images in a given preparation), its image was replaced by the nearest intact sister section. Images were aligned using the Align tool, including automatic alignment supplemented by manual fitting. Once aligned, signal was annotated using a manually set threshold. Surface files were made of label files that were resampled to a voxel-size of approximately 10×10×10 μm.

## Results

### The adult heart of monitors has a left and a right ventricle

The monitor ventricle consists mostly of trabeculated myocardium, as is the case in other squamates (Fig. 1). In the adult monitor ventricle (*V. exanthematicus* and *V. salvator*), the trabeculations were organized into a thick-walled and left-sided systemic ‘left ventricle’ and a thin-walled and right-sided pulmonary ‘right ventricle’, or cavum pulmonale (Fig. 2A, Movie S1). In the transverse plane, the ‘left ventricle’ was circular and the ‘right ventricle’ formed a crescent at its right side (Fig. 2B, Movie S2). The two ventricles were separated by the muscular ridge and the bulbuslamelle (Fig. 2B). In the apical half of the ventricle, the two septa were fused and appeared as one septum. We noticed highly echogenic lamellae in the inner-most part of the cavum venosum. These lamellae correspond to the so-called bulboauricular lamellae of Greil (Greil, 1903; B. Jensen et al., 2013). In a large heart of *Varanus salvator*, these lamellae were thick and constituted the inner-most part of the muscular ridge and bulbuslamelle (Fig. 2C). They extended from the atrioventricular canal to the point where the muscular ridge and bulbuslamelle abut during contraction.

**Fig. 2.**
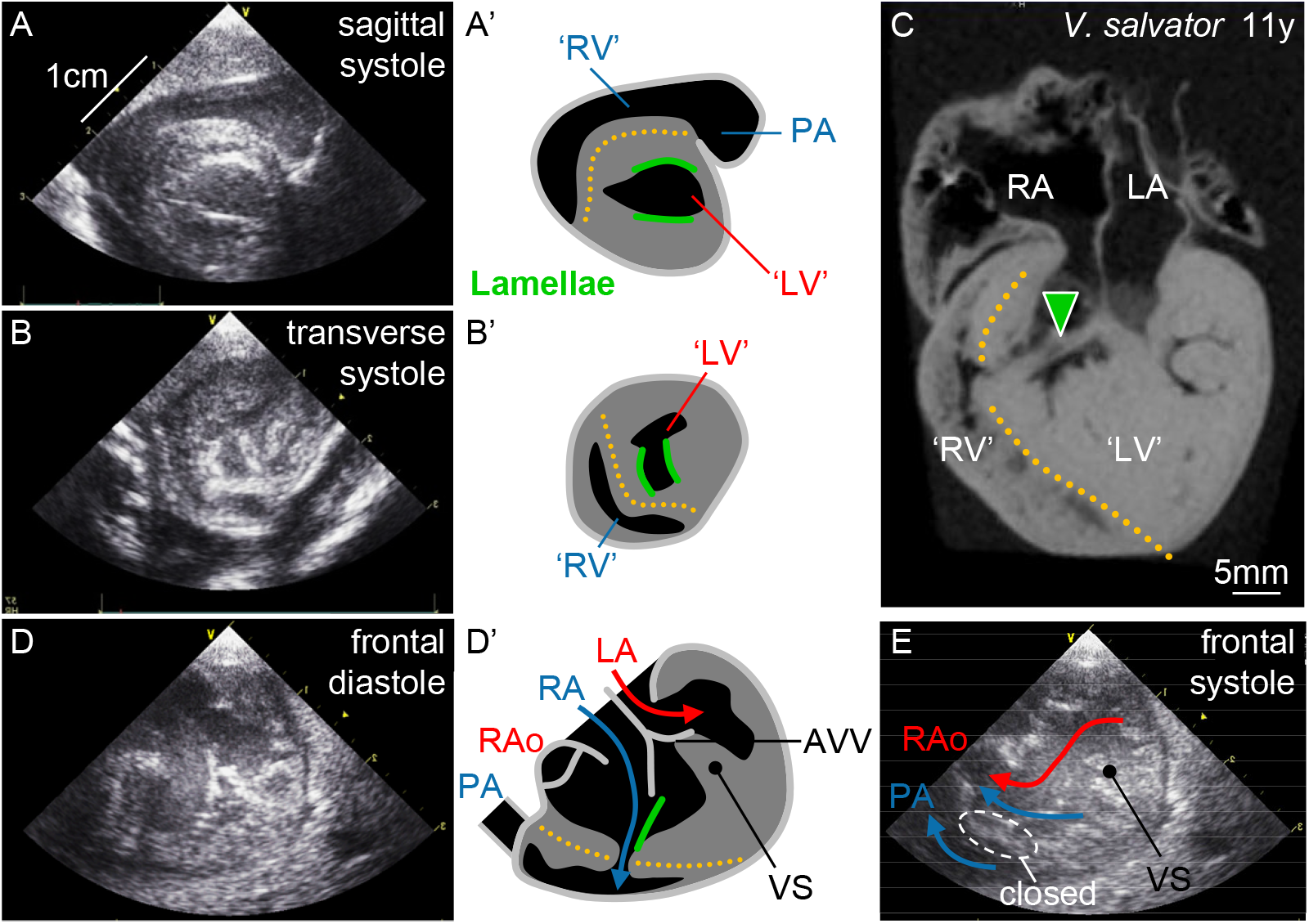
The monitor ventricle is divided. **A-A’**. Echocardiography of the ventricle of the Savannah monitor showing a densely muscular left ventricle (‘LV’) and a less muscular right ventricle (‘RV’) leading to the pulmonary artery (PA). The echogenic lamellae (green) are the bulboauricular lamellae of Greil (Greil, 1903). They constitute the inner-most part of the muscular ridge and bulbuslamelle. **B-B’**. In the transverse plane, the right ventricle forms a semi-circle as it does in mammals and birds, nestled against the contracting muscular ridge and bulbuslamelle (orange dotted line). **C**. The contracted ventricle of the Water Monitor (visualized with MRI) is similar to the ventricle the Savannah monitor and shows a pronounced bulboauricular lamella (green arrowhead). **D-E**. In ventricular diastole of the Savannah Monitor (**D**), the atrioventricular valve (AVV) contacts the vertical septum (VS) and the blood streams of the left atrium (LA) and right atrium (RA) are separated along it, as the atrial blood streams are along the ventricular septum of mammals and archosaurs. To reach the right ventricle, blood from the right atrium has to pass through the left ventricle and the small gap between the muscular ridge and bulbuslamelle (blue arrow). In systole (**E**), contractions close the gap (“closed”) and the muscular ridge and bulbuslamelle now separates the low pressure right ventricle from high pressure left ventricle, as does the ventricular septum of mammals and archosaurs (**D-E** are screenshots of a previously published movie (B. Jensen et al., 2014)). RAo, right aorta.

In diastole, the atrioventricular valve achieved direct contact to a prominent vertical septum (Fig. 2D, Movie S3). The vertical septum was positioned entirely within the cavity of the left ventricle (the cavum arteriosum and cavum venosum combined). Both atrial blood flows were therefore received in the left ventricle. As atrial blood entered the ventricle, the space between the muscular ridge and the bulbuslamelle widened, allowing the right atrial blood to reach the cavum pulmonale (the right ventricle). In systole, this space closed again as the muscular ridge and the bulbuslamelle abutted. Then, the ‘right ventricle’ ejected into the pulmonary trunk and the ‘left ventricle’ ejected into the aortae (Fig. 2E, Movie S3). Septal structures of the monitor ventricle showed resemblance both functionally and anatomically to ventricular structures of mammals and archosaurs, and therefore we wanted to know next whether the septal structures of monitors develop as in the four-chambered heart.

### Morphogenesis of the monitor heart

The development of the morphology of the monitor heart was predominantly studied in the Mangrove monitor series, where all hearts were stained for cardiac muscle. In the earliest specimen, from 10 days post oviposition, the heart was much like the chicken heart around 4 days of development, that is, myocardium was detected in the sinus venosus, sinuatrial junction, atria, atrioventricular canal, ventricle, and outflow tract (Fig. 3A-B). The ventricle was relatively small, whereas the atrioventricular canal and myocardial outflow tract, the conus arteriosus, were proportionally large, showing that the heart was immature (Fig. S1). In the atria, trabeculation was just beginning to appear and the atrial septum was a low myocardial ridge with a mesenchymal cap in the atrial roof (Fig. 3A). A non-muscular dorsal mesenchymal protrusion was present in the dorsal wall of the atrium. The atrioventricular canal was circular in cross-section and its interior was occupied by the large dorsal and ventral mesenchymal cushion (Fig. 3A). Its position was to the immediate left of the body midline as given by the position of the notochord, foregut and trachea. Trabeculation of the ventricle was prominent in the outer curvature and absent in the inner curvature (Fig. 3A). Distally, the trabeculated muscle was sponge-like, while proximally it was organized into some 12 sheets that were oriented approximately dorso-ventrally. The outflow tract was proportionally long, it was myocardial to the vicinity of the pericardial reflection, and its interior was dominated by mesenchyme but the lumen was not divided (Fig. 3A).

**Fig. 3.**
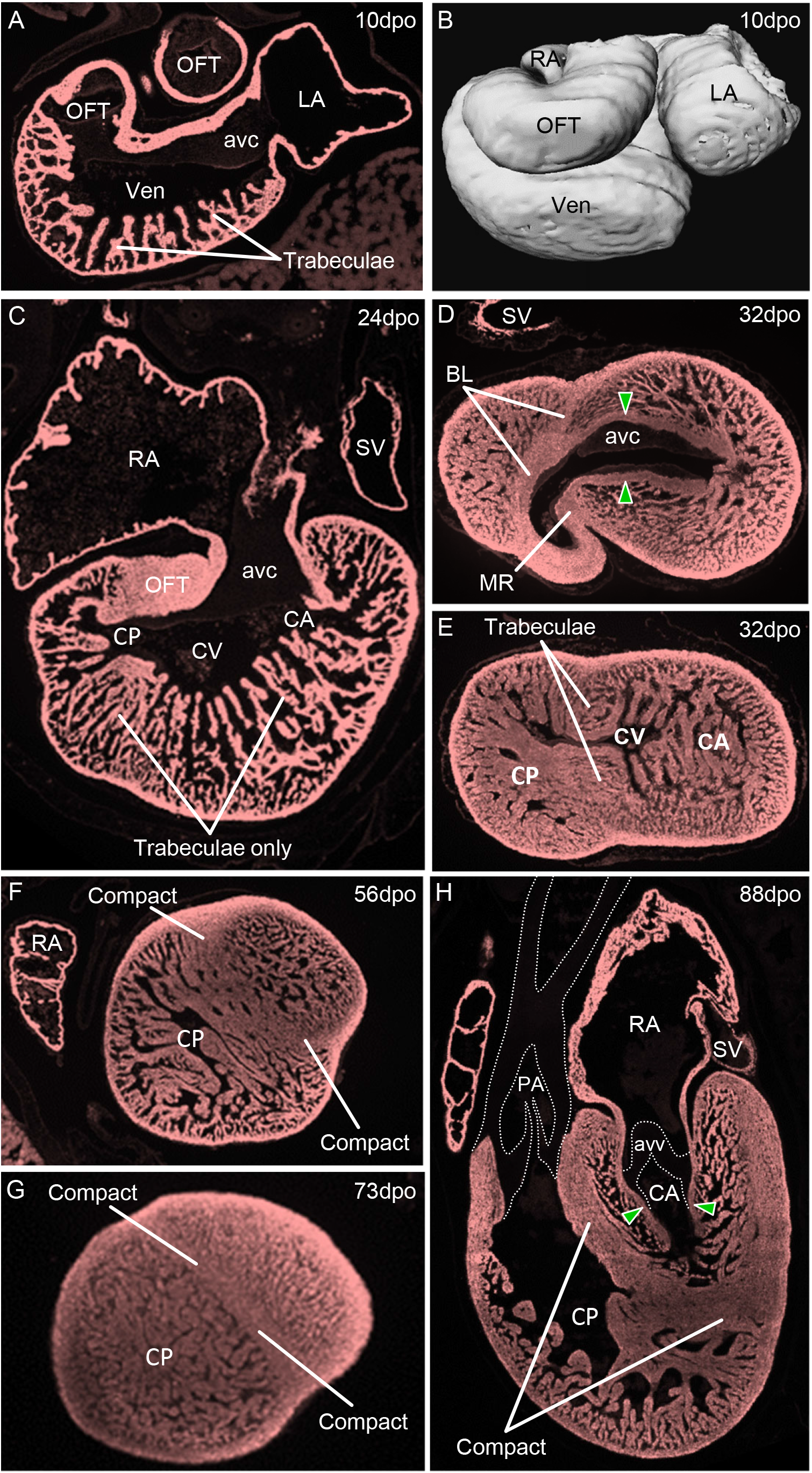
Morphogenesis of the Mangrove monitor heart. **A**. Frontal section of the heart of the specimen of 10 days post oviposition (dpo), showing myocardium in all compartments of the heart (sinus venosus is not seen), and the trabeculae of the outer curvature of the ventricle (Ven). **B**. 3D model of the 10 dpo heart, showing the long outflow tract (OFT). **C**. At 24 dpo, ventricular trabeculae are extensive. D. At 32 dpo, the muscular ridge (MR) and the bulbuslamelle (BL) are recognizable by compact myocardium. The bulboauricular lamellae (green arrowheads) are much developed. **E**. At mid-height of the ventricle, the compact myocardium of the muscular ridge and the bulbuslamelle blends into trabeculae. **F-G**. At 56 dpo, some ventricular trabeculae of the apical region show a more compact organization and at 73 dpo the apex is divided by a compact septum. **H**. At 88 dpo, compact myocardium fully separates the cavum arteriosum (CA) from the cavum pulmonale (CP). The bulboauricular lamellae (green arrowheads) extend deep into the cavum arteriosum. avc, atrioventricular cushion; avv, atrioventricular valve; CV, cavum venosusm; LA, left atrium; PA, pulmonary artery; RA, right atrium; SV, sinus venosus.

In the next investigated age, 24 days post oviposition (Fig. 3C), all three sinus horns were distinct vessels (the left, right, and posterior sinus horn) and their myocardium extended to the pericardial reflection. The atrial septum was so developed that the primary ostium was almost closed and secondary foramina had formed. The atrioventricular canal remained to the left of the body midline while its cushions had begun to merge. Ventricular trabeculation was extensive. While the formation of the muscular ridge had begun near the base of the outflow tract, the outer curvature of the ventricle comprised trabecular myocardium only (Fig. 3C). The outflow tract was myocardial only in the proximal half, the distal half being arterial, and the lumen was divided by mesenchyme into three channels, those of the future pulmonary artery and the left and right aorta. At this age, the proportions of the principal compartments of the heart were almost like in the formed heart, only the ventricle still had to increase a bit at the expense of the regression of the myocardial outflow tract (Fig. S1).

By 32 days post oviposition, the primary ostium was closed and spots of myocardium were detected within the dorsal mesenchymal protrusion. In the base of the ventricle, both the muscular ridge and the bulbuslamelle were characterized by a core of compact myocardium (Fig. 3D). Towards the apex, the compact myocardium dissolved into trabeculae (Fig. 3E). The trabecular sheets beneath the atrioventricular cushions were of approximately equal height and there was no obvious vertical septum at this age. The bulboauricular lamellae were becoming prominent (Fig. 3D-E). In the next investigated ages, 56 and 73 days post oviposition, parts of the apical trabeculae started to coalesce to a compact organization (Fig. 3F-G). At 73 days post oviposition, the lumen of coronary vessels could be identified by a high density of SYTOX blue (the erythrocytes are nucleated, data not shown). These were sub-epicardial only, predominantly where the muscular ridge and the bulbuslamelle made contact with the outer shell of compact muscle, and we did not detect coronary vessels within the compact of any septa. At 88 and 144 days post oviposition, the atrioventricular canal remained to the left of the aortic base. Compact muscle now enclosed the cavum arteriosum and venosum and thus separated these parts from the cavum pulmonale (Fig. 3H) (except at the gap between the muscular ridge and the bulbuslamelle). Coronary vessels were found in the outer shell of ventricular compact muscle and they originated from an approximately 50μm wide orifices in the dorsal valve sinus of the left aorta. We could not detect coronary vessels within the compact of the septa. The bulboauricular lamellae were prominent (Fig. 3H). Much like in the embryonic chicken ventricle, the monitor ventricle develops extensive trabeculae followed by septation by compaction, so we hypothesized that the monitor septa would express genes associated with the development of the full ventricular septum.

### Monitor septal structures express orthologues of transcription factors of the full ventricular septum

For *in situ* hybridization of embryonic lizards, we focused on the stages when septal structures have formed and when the ventricle has grown to approximately the proportion of the total cardiac mass that is seen in adult animals, i.e. some 70-80% (Fig. S1); in *Varanus acanthurus* this corresponded to ca. 35 days post oviposition and in *Norops sagrei* to Sanger stage 11 (B. Jensen et al., 2013; Sanger et al., 2008). In the monitors, *Tbx3* was expressed in the bulboauricular lamellae from the atrioventricular canal to the vicinity of the vertical septum (Fig. 4A). This expression, and the bulboauricular lamellae, extended deeper into the ventricle than it did in *Norops* (Fig. 4A-B). In the monitors, *Tbx5* was expressed throughout the ventricle, with no gradient between the left and right sides (Fig. 4C). The myocardial outflow tract did not express *Tbx5*. Within the ventricle, Tbx5 was particularly rich in the bulboauricular lamellae and therefore enriched near the vertical septum and in the innermost parts of the muscular ridge and the bulbuslamelle (Fig. 4C). In *Norops*, the bulboauricular lamellae were less developed and they were not enriched in *Tbx5* as in the monitors (Fig. 4D). In the monitors, *Irx1* and *Irx2* were expressed near the vertical septum and in the innermost parts of the muscular ridge and the bulbuslamelle (Fig. 4E, G). *Irx1* and *Irx2* were similarly expressed in the ventricles of the monitors and *Norops*, although the patterns were more pronounced in the monitors (Fig. 4E-G). Figure 4H shows a 3D reconstruction of the myocardium in the *Varanus indicus* ventricle (56 dpo) that expressed *Irx1, Irx2*, and *Tbx5*. In *Norops, Irx1* expression extended beyond the domain of *Tbx5* expression and into the outflow tract myocardium (Fig. 4F, S2). Figures 4I-J show *Irx1* and *Irx2* expression in the septal region of chicken and mouse, respectively. Outside the heart, *Tbx3, Tbx5, Irx1*, and *Irx2* were also detected in the developing trachea and bronchi (Fig. 4). Taken together, the expression of genes associated with the ventricular septum of mammals and birds was found around the ventricular septal structures of monitors. The myocardium immediately next to the vertical septum of the monitors had a molecular phenotype comparable to the atrioventricular bundle of mammals and birds, which led us to hypothesize that this myocardium would be activated early, as it is in mammals and birds.

**Fig. 4.**
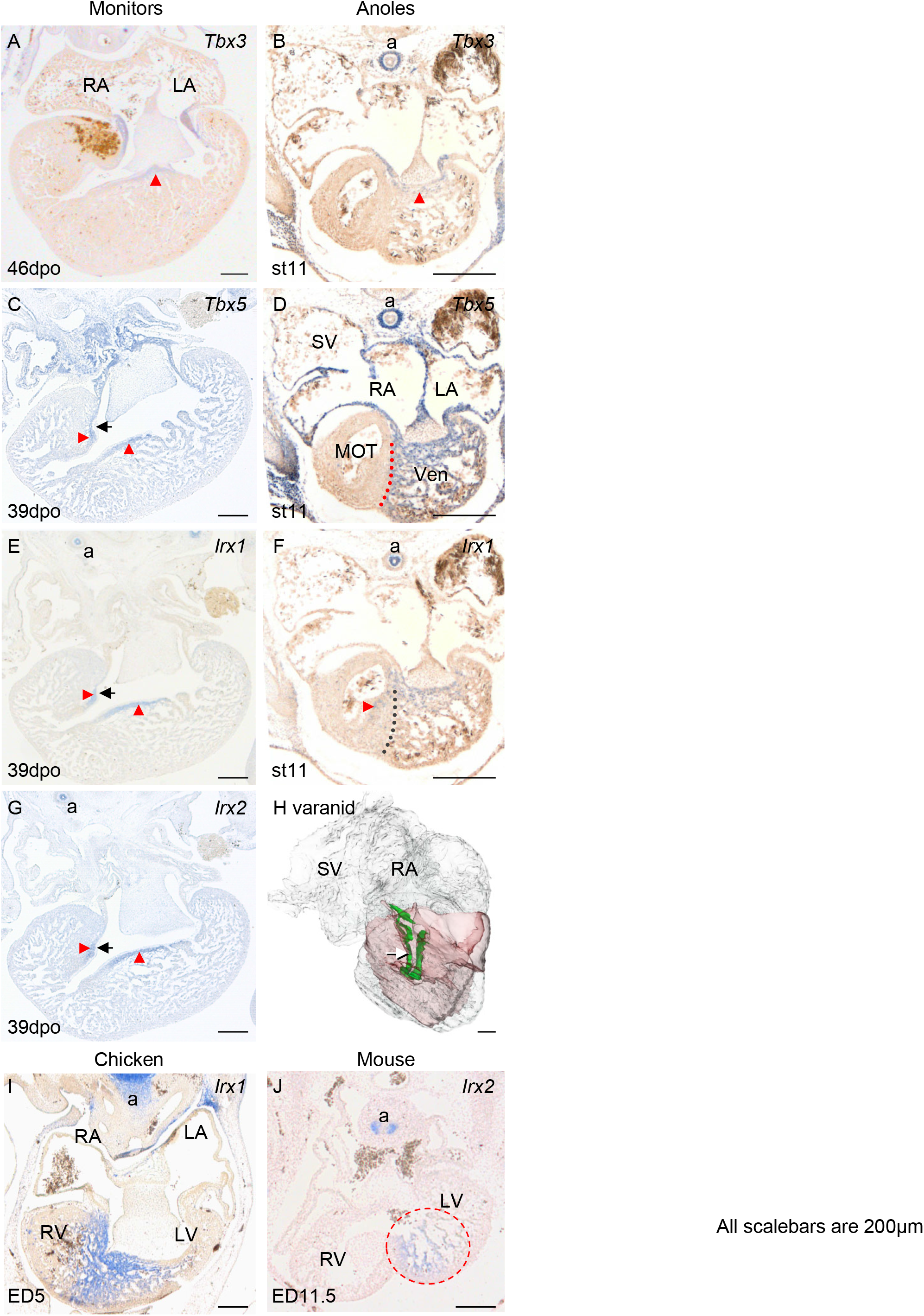
Septal structures in monitors, anoles, chicken, and mouse express homologous genes. **A-B**. In monitors, *Tbx3* was expressed from the atrioventricular canal to the vicinity of the vertical septum (red arrowhead), much like in the anole, but more pronounced. **C**. *Tbx5* was expressed throughout the monitor ventricle and was particularly rich in the bulboauricular lamellae and the connected crests of trabecular sheets (red arrowheads). The black arrow points to outflow tract cushion on the bulbuslamelle. **D**. In the Brown Anole, *Tbx5* expression was uniform in the ventricle, but absent from the myocardial outflow tract (MOT). **E-G**. In monitors, the bulboauricular lamellae and the connected crests of trabecular sheets expressed *Irx1* and *Irx2* (red arrowheads), and in the Brown anole (**F**), *Irx1* was expressed like in the monitors and extended beyond the *Tbx5* gradient (dotted line) to the muscular ridge. **H**. Green color shows the myocardium of monitors that was rich in *Tbx3, Tbx5, Irx1*, and *Irx2* (green), together with the totality of the muscular ridge and bulbuslamelle (transparent red) in a reconstruction of all myocardium (transparent grey) of a 56dpo Mangrove Monitor seen from the right. The arrow indicates the gap between the muscular ridge and bulbuslamelle. **I**. Ventricular expression of *Irx1* in the forming septum of chicken. **J**. Ventricular expression of *Irx2* in the forming septum of mouse (encircled). Notice that transcripts were detected in the developing airways; a in B, D-G, I-J. LA, left atrium; LV, left ventricle; RA, right atrium; RV, right ventricle; SV sinus venosus. Scale bars 200 μm.

### Early activation of the ventricular myocardium in the vicinity of the vertical septum in monitors

Optical mapping of cardiac action potentials was performed on 6 Ridge-tailed monitor embryos ranging from days 21 to 46 post oviposition, spanning the period when ventricular septa are formed. Despite substantial growth of the embryo and heart during this period, the intrinsic heart rate, atrial activation time, atrioventricular delay, and ventricular activation times were essentially constant (Fig. 3S). The left side of the ventricle was consistently activated earlier than the right side (Fig. 5A). On the left side, the earliest epicardial activation was always observed in the vicinity of the forming vertical septum (Fig. 5A). In the oldest embryos, this was seen as a site of activation relatively deep in the ventricle (Fig. 5C), which was different from the primitive base-to-apex activation of even older Leopard Geckos (Fig. 5B). This pattern of activation could be elicited by stimulation in the apical region of the left chamber (only performed in the oldest and largest specimens, data not shown). Leopard Geckos showed the primitive base-to-apex activation pattern throughout development (N=11, Fig. 5C-D).

**Fig. 5.**
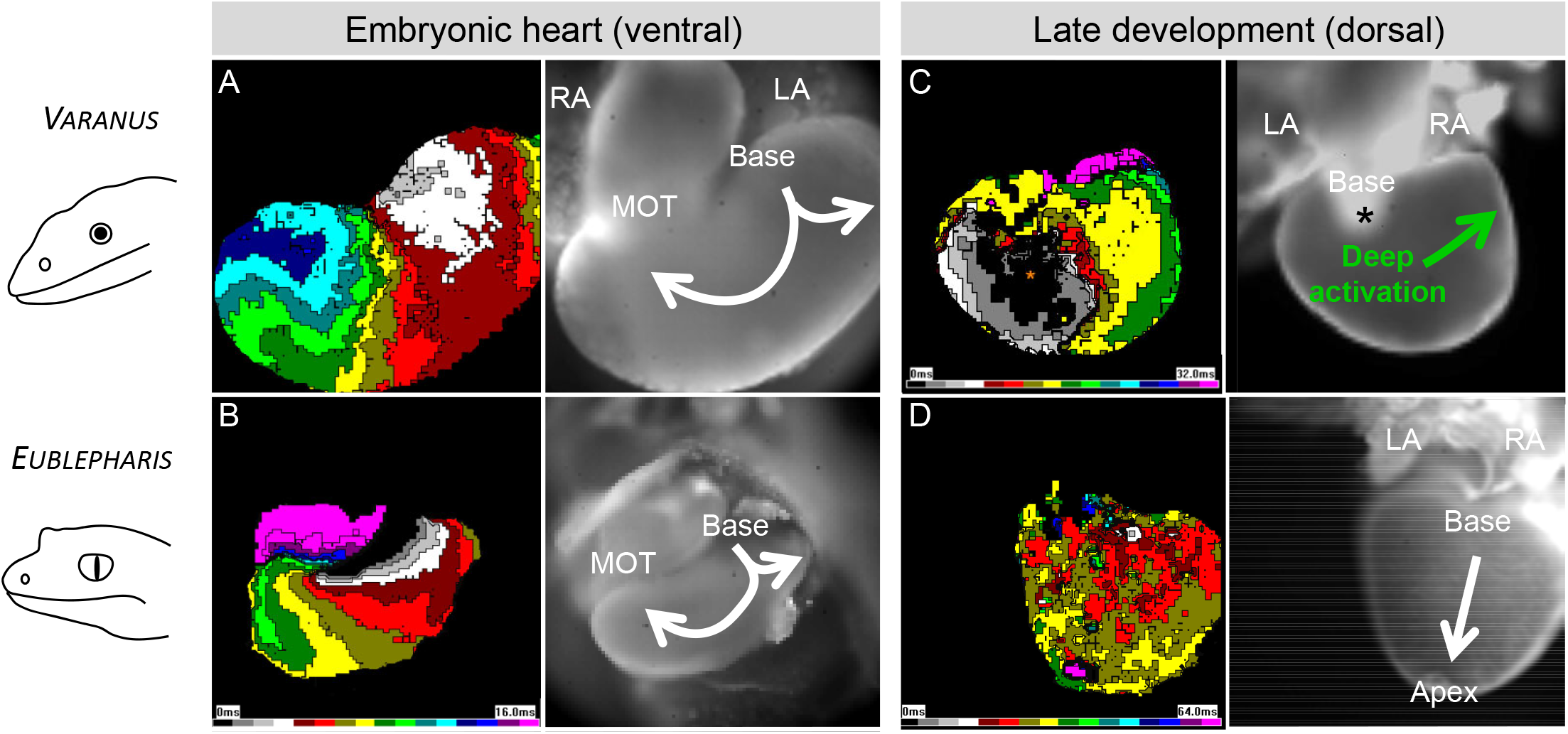
Ventricular activation becomes specialized in monitors. **A-B**. In the embryonic hearts of both Ridge-tailed Monitor (21dpo) and Leopard Gecko (7 dpo), ventricular activation initiated in the base and swept over the apex and towards the myocardial outflow tract (MOT). **C**. In later development of the monitors (46dpo shown), early activation was deep (black field, asterisk) towards the apex on the left and therefore specialized. The black asterisk indicates the approximate position of the crest of the vertical septum. **D**. In later development of the Leopard Gecko (53 dpo), the earliest activation was in the ventricular base, and its spread exhibited the primitive base-to-apex pattern. LA, left atrium; RA, right atrium.

### Expression of Myh6 suggests the bulboauricular lamellae are exceptionally developed in monitors

We noticed in the monitors that *Myh6* was readily detected in the atria, atrioventricular canal and the innermost ventricular myocardium - in particular in the bulboauricular lamellae (Fig. S4). In the earliest investigated embryo of *Norops, Myh6* was readily detected in the atria, the atrioventricular canal, and sinus venosus, whereas the signal in the ventricle was very faint and broad (Fig. S1). No signal was detected in the ventricle of later stages (Fig. S1). In embryonic mammals and chicken, *Myh6* is initially expressed in the ventricle but becomes confined to the forming atrioventricular bundle (de Groot et al., 1987; A. F. M. Moorman et al., 1998; Yutzey, Rhee, & Bader, 1994). Across a phylogenetically diverse set of non-crocodylian reptiles, *Myh6* was detected in the atria and atrioventricular canal of all species (Fig. 6). Only in monitor lizards, however, we observed *Myh6* in the ventricle as well (Fig. 6). This expression was prominent in the bulboauricular lamellae, which were much more developed than in the other reptiles (Fig. 6). In the Leopard Geckos, used for optical mapping, *Myh6* was expressed in a pattern like in the Anoles, *i.e*. readily detected in the atria (and sinus venosus), but not in the ventricle (Fig. 6). These observations corroborated the hypothesis that there is an association between the pronounced septation, the developmental state of the bulboauricular lamellae, and a specialized manner of ventricular electrical activation.

**Fig. 6.**
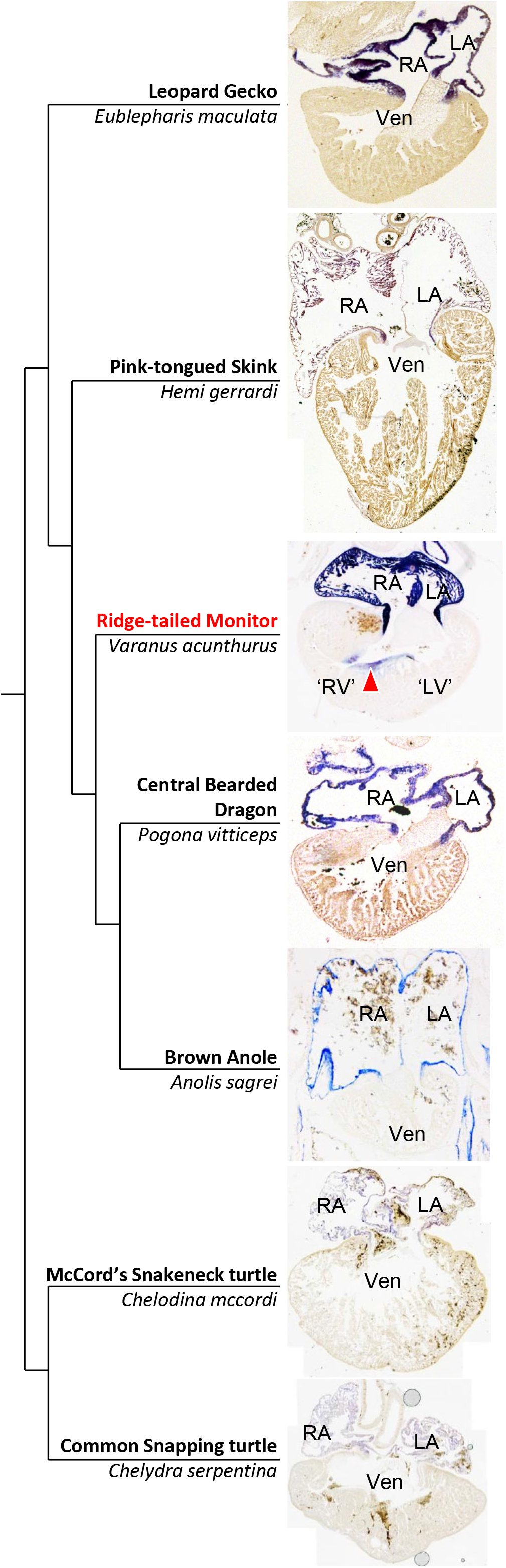
Cardiac expression of *Myh6* identifies the bulboauricular lamellae in monitors only. LA, left atrium; ‘LV’, monitor left ventricle; RA, right atrium; ‘RV’, monitor right ventricle; Ven, single ventricle.

## Discussion

Monitor lizards are the only known lizards with a functionally divided ventricle. Here we established that the dividing septa express *Irx1, Irx2, Tbx3*, and *Tbx5*. In all studied ventricles with hemodynamically distinct left and right sides, *i.e*. human, mouse, chicken, and crocodylians, these transcription factors are expressed in the septal structures (Bruneau et al., 1999; Christoffels, Keijser, et al., 2000; Hoogaars et al., 2004; B. Jensen et al., 2018; Sizarov et al., 2011; Yamada et al., 2000). Homologous molecular mechanisms, therefore, seem to underlie the processes of ventricular septation in all these instances. Our results suggest that the various degrees of septation encountered in Sauropsida (reptiles and birds) and Synapsida (mammals) are variations of the same building plan inherited from their common amniote ancestor. Incidentally, we found the same transcription factors to be expressed in the trachea and bronchi, as they are in the developing trachea and bronchi of mouse and chicken (Chapman et al., 1996; Christoffels, Keijser, et al., 2000; Gibson-Brown, Agulnik, Silver, & Papaioannou, 1998). Therefore, homologous molecular mechanisms also appear to drive the formation of the main airways despite the considerable anatomical differences in the formed respiratory systems of reptiles, birds, and mammals (Farmer & Sanders, 2010; Perry & Sander, 2004; Schachner, Cieri, Butler, & Farmer, 2014; Steimle et al., 2018).

The muscular part of the human ventricular septum has been described by anatomists as having distinct components, specifically an inlet part (dorsal/posterior and a-trabecular), an apical part (surrounded by the trabeculae), and an outlet part (ventral/anterior and a-trabecular) (Robert Henry Anderson & Becker, 1980; Van Mierop & Kutsche, 1985; Wenink, 1981). Molecular markers that distinguish these components remain to be found. From comparative anatomical studies, it was concluded that the outlet part corresponds to the muscular ridge (Benninghoff, 1933; R.E. Poelmann et al., 2014; Van Mierop & Kutsche, 1985; Grahame J.W. Webb, 1979). The muscular ridge of non-crocodylian reptiles, including monitors, has a gradient of *Tbx5*, as does the ventricular septum of mammals, chicken, and crocodylians (B. Jensen et al., 2018; Koshiba-Takeuchi et al., 2009; R.E. Poelmann et al., 2014). However, the muscular ridge is not positioned beneath the atrioventricular junction as is the full ventricular septum and the prevailing view is that the muscular ridge cannot be equated wholesale to the muscular part of the full ventricular septum (Benninghoff, 1933; R.E. Poelmann et al., 2014; Van Mierop & Kutsche, 1985; Grahame J.W. Webb, 1979). A gradient of *Tbx5* is also present on the ventral-most part of the bulbuslamelle, but in mammals and birds this part seems to contribute to the right atrioventricular valve rather than the septum (B. Jensen & Moorman, 2016; B. Jensen et al., 2013; Lamers, Viragh, Wessels, Moorman, & Anderson, 1995). The dorsal-most part of the bulbuslamelle is thought to contribute to the ventricular septum (Benninghoff, 1933; Greil, 1903; B. Jensen et al., 2014; G.J.W. Webb, 1979), but this part was not enriched for any of the investigated transcripts and did not have a gradient of *Tbx5*. Therefore, when comparing the muscular ridge and the bulbuslamelle of monitors to the ventricular septum of mammals and birds, there is a shared expression of key transcription factors but there is a divergence in the anatomy of the myocardial tissues that express these genes. The vertical septum is beneath the atrioventricular junction and the myocardium next to it is enriched in transcripts associated with the atrioventricular bundle of mammals and archosaurs (Bakker et al., 2008; B. Jensen et al.). The early electrical activation near the vertical septum corroborates this interpretation.

The muscular ridge and the bulbuslamelle could both be identified by the compact organization of their myocardium. In the early stages, this myocardium blended into apical trabeculae while at later stages the compact muscle of the muscular ridge and the bulbuslamelle could be followed to the apex. This strongly suggests that some trabeculae had undergone compaction. Compaction is considered a key process of the formation of the ventricular walls and septum of mammals and birds (Captur, Syrris, Obianyo, Limongelli, & Moon, 2015; Sedmera, Pexieder, Vuillemin, Thompson, & Anderson, 2000), although the importance of the process may vary between species and its contribution to pathology is debated (R. H. Anderson et al., 2017; Finsterer, Stollberger, & Towbin, 2017).

The final component of the full ventricular septum is the membranous septum (Cook et al., 2017; Greil, 1903; B. Jensen et al., 2014; G.J.W. Webb, 1979). It may have precursors in reptiles, since large cushions of connective tissue can be found in monitors and pythons where their muscular ridge and bulbuslamelle abut (B. Jensen et al., 2014; Bjarke Jensen, Nyengaard, Pedersen, & Wang, 2010). If these cushions were to fuse, not only would the ventricle be fully divided, but the membranous septum would also be closely aligned with the arterial wall that separates the pulmonary artery from the aortae.

In developing mammals and archosaurs, the atrioventricular canal undergoes a pronounced right-ward expansion during ventricular septation whereby the right atrium remains in contact with the right ventricle (B. Jensen & Moorman; B. Jensen et al.; Lamers et al.). This process includes the formation of the right atrioventricular valve and failed expansion results in congenital malformations such as tricuspid atresia and double inlet left ventricle that have been reported in multiple mammalian species (B. Jensen & Moorman, 2016; Michaëlsson & Ho, 2000). In squamate reptiles with an undivided ventricle, the atrioventricular canal is positioned to the left of the body midline (B. Jensen et al., 2013) and does not undergo a rightward expansion (B. Jensen & Moorman, 2016). It is then remarkable that the atrioventricular canal of monitors does not undergo the rightward expansion despite the functional division of the ventricle. One consequence is that the cavum venosum of monitors serves as the conduit for systemic venous blood to the cavum pulmonale in ventricular diastole and serves as the conduit for pulmonary venous blood in systole (Fig. 2D-E). Anatomists and physiologists have long emphasized that the heart of monitors can be seen as a conceptual stage between the hearts of non-crocodylian reptiles and the fully septated hearts of crocodylians, birds, and mammals (Acolat, 1943; Brücke, 1852; Greil, 1903; B. Jensen et al., 2014). The primitive state of lack of right-ward expansion of the atrioventricular canal concomitant with the specialized state of deep left ventricular activation corroborates this notion.

Comparing lizards, crocodylians, birds, and mammals, we find a considerable overlap in the functional anatomy, electrophysiology, and molecular phenotype of the septation of the ventricles of amniotes. Our results suggest that Sauropsida and Synapsida inherited precursor structures from their common amniote ancestor. Particular features evolved within each lineage (Cook et al., 2017), but we propose they could be seen as variations on a common design.

## Acknowledgements

BJ was supported by The Carlsberg Foundation [CF14-0934]. DS was funded by Ministry of Education [PROGRESS Q38], Czech Academy of Sciences [RVO: 67985823], Grant Agency of the Czech Republic [16-02972S]. TW was supported by the Danish Natural Science Research Council.

---

**Fig. S1.**
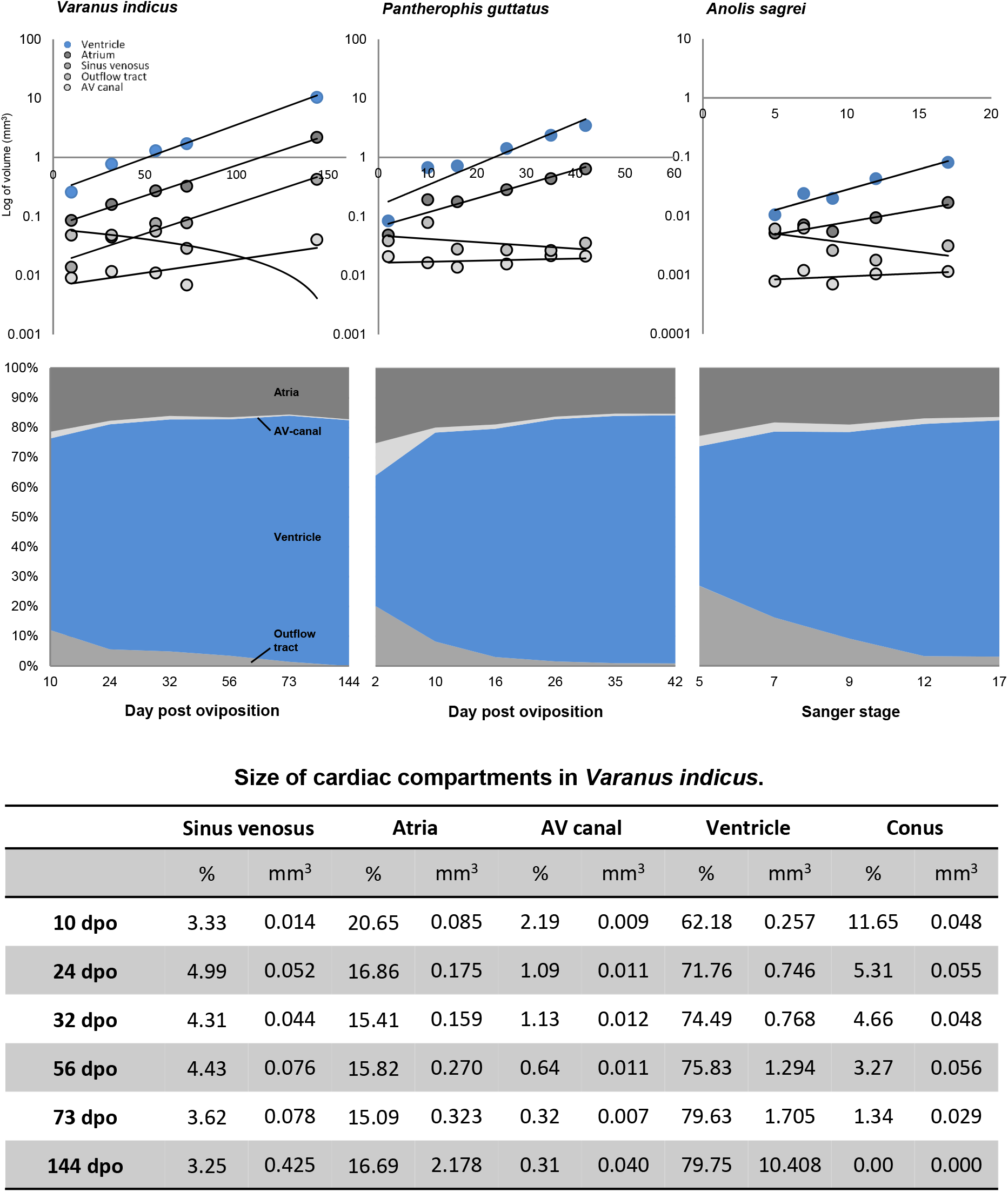
Growth of the cardiac compartments of Mangrove Monitor resembled the growth seen in the Brown Anole and the Corn Snake. In Mangrove Monitor, the ventricle, the atria, and the sinus venosus exhibited similar rates of exponential growth. By the end of the studied developmental window, the monitor ventricle constituted a similar proportion of the total cardiac mass as in Corn Snake and Brown Anole. The data for Corn Snake and Brown Anole is adapted from (B. Jensen et al., 2013).

**Fig. S2.**
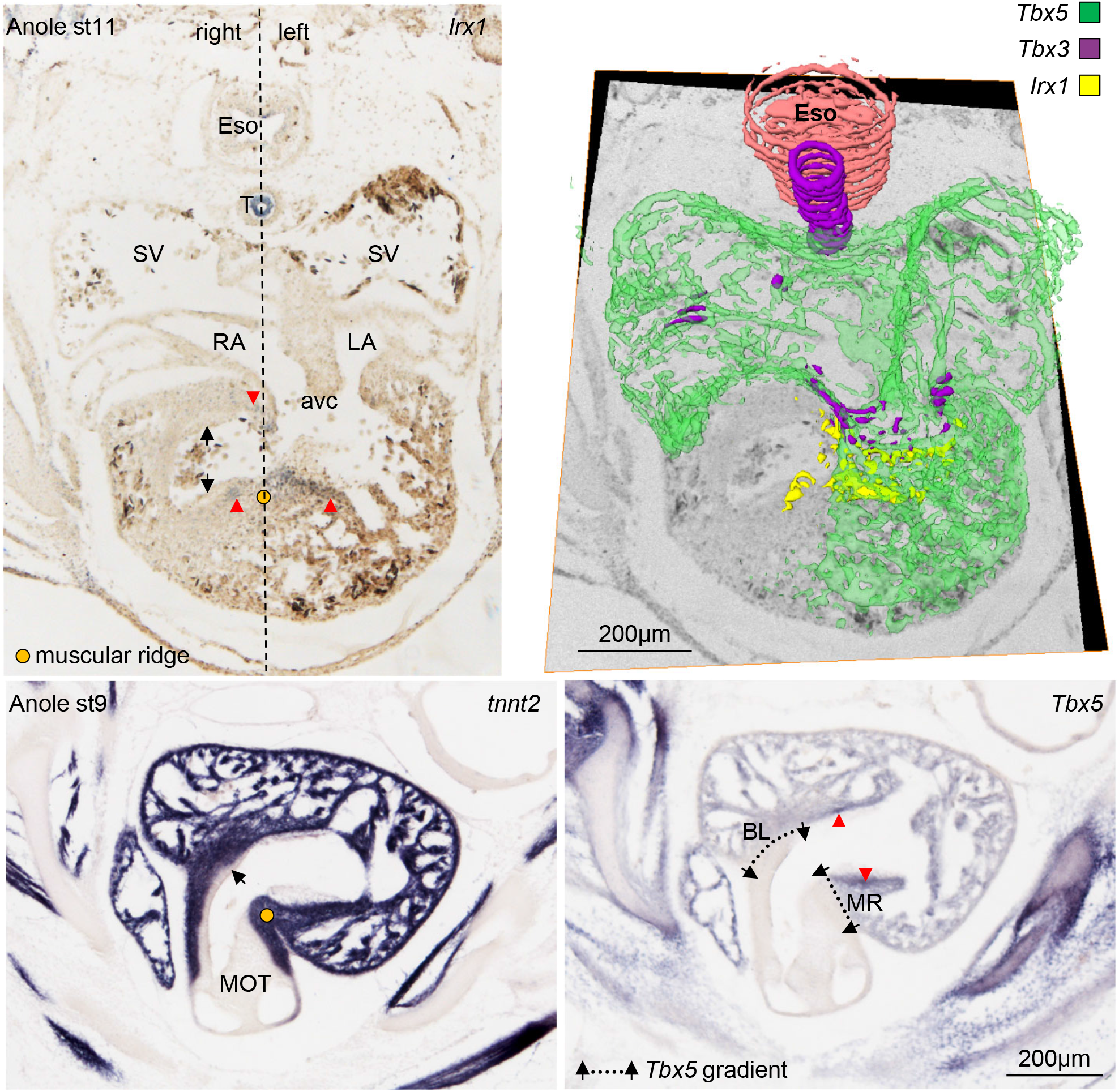
Reconstruction of the expression pattern of *Tbx5, Tbx3*, and *Irx1* in the Brown Anole. Expression of *Irx1* (red arrows) extends to the right of the body midline and into the myocardium of the myocardial outflow tract (black arrows point to outflow tract cushions) on the right side of the body midline. These parts correspond to the inner-most parts of the muscular ridge and bulbuslamelle of the monitors. In the stage 9 anole, ventricular expression of *Tbx5* is strongest near the atrioventricular canal (red arrowheads), whereas gradients between no expression and expression can be found on the muscular ridge (MR) and bulbuslamelle (BL). avc, atrioventricular canal; Eso, esophagus; LA, left atrium; MOT, myocardial outflow tract; RA, right atrium; SV, sinus venosus; T, trachea.

**Fig. S3.**
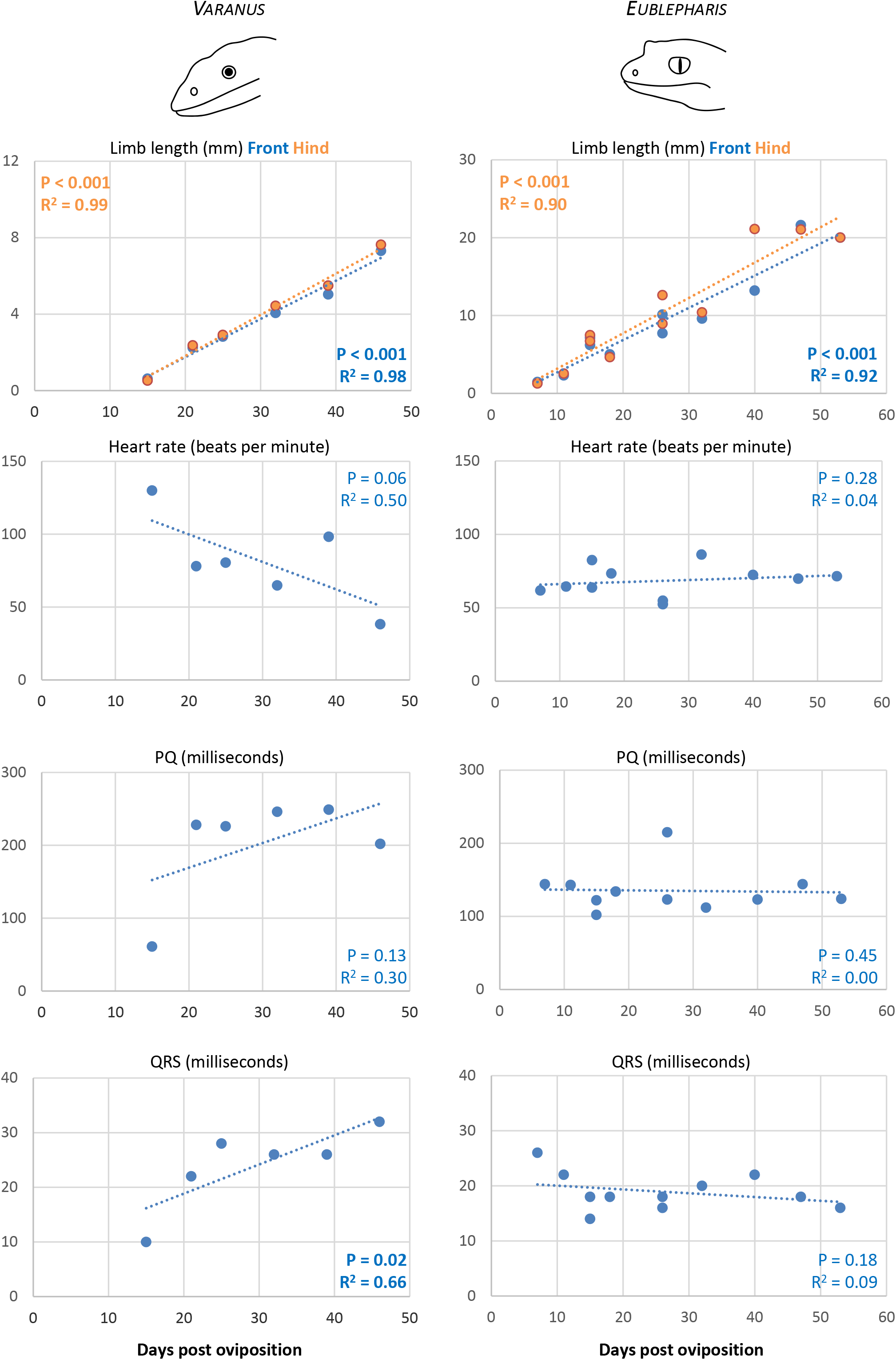
Growth and electrophysiological parameters in development of the Ridge-tailed Monitor and the Leopard Gecko. The animals had a steady increment in size throughout the developmental period of septum formation as measured by the length of the right front limb (Front) and right hind limb (Hind). In the graphs are the values of the Pearson correlation tests (P) and R^2^ of linear regressions (R^2^). In the same period, the electrophysiological parameters were mostly stable, as has been reported for the intermediate and late developmental stages of other species of reptiles and birds (Gregorovicova et al., 2018). High heart rate associates with shorter durations of the PQ and QRS, and the high heart rates of the youngest monitor accounts for most of the apparent differences between the Ridge-tailed Monitor and the Leopard Gecko.

**Fig. S4.**
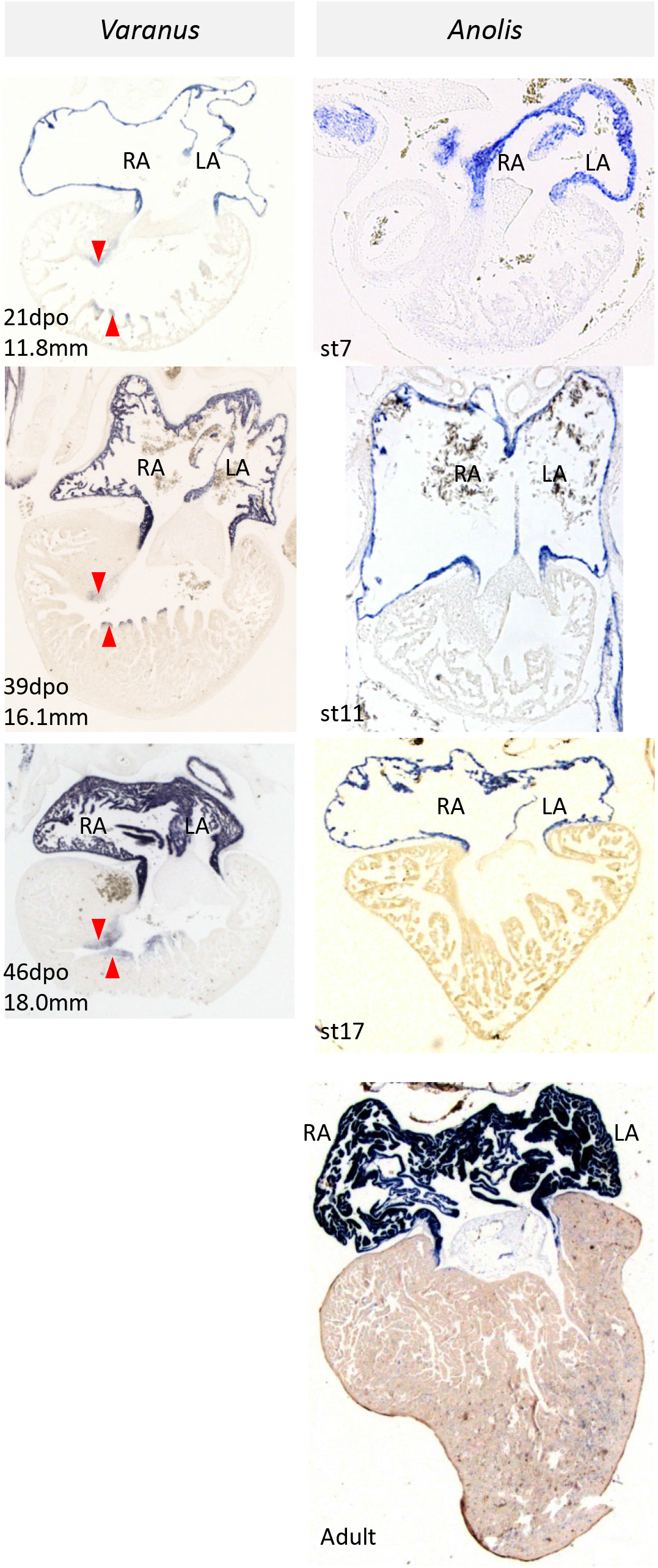
Expression of *Norops* atrial myosin heavy chain (*Myh6*) in development of the Ridge-tailed Monitor and Brown Anole. In the monitors, *Myh6* (red arrowheads) was expressed in a subset of myocytes in the ventricle of all investigated stages (21-46dpo, 11.8-18.0mm crown-rump length), whereas the anole lizards never showed similar expression (Sanger stage 7 to adult). LA, left atrium; RA, right.

**Movie S1**. Echocardiography of an anaesthetized Savannah Monitor in a sagittal-like plane showing the systemic, or ‘left’, ventricle as dense and circular, with the pulmonary, or ‘right’, ventricle nestled around it.

**Movie S2**. Echocardiography of an anaesthetized Savannah Monitor in the transverse plane showing the ‘left’ ventricle as dense and circular with the ‘right’ ventricle nestled around it.

**Movie S3**. Echocardiography of an anaesthetized Savannah Monitor in the frontal plane showing the atrioventricular valves interacting with a prominent ridge, the vertical septum, deep within the ‘left’ ventricle.

